# Towards a Physiological Scaling Law: Model Quality vs. Cohort Size for Stochastic Sequence Data

**DOI:** 10.64898/2026.08.11.744303

**Authors:** Gayathri Sunil, Bhuvana Ravi Kumar, Bharath Ramsundar, Sandya Subramanian

**Affiliations:** University of Massachusetts Amherst; University of California Davis; Deep Forest Sciences, Inc.; University of California Berkeley

## Abstract

Scaling laws help determine the optimal data size for training large models but are established in domains where the target is deterministic. Physiological signals are different: heartbeat sequences are stochastic, so part of the error is irreducible even with large amounts of data. Metrics such as MAE do not account for non-deterministic behavior, and therefore assessing scaling requires evaluating distributional calibration (measuring how well predicted probability densities capture true conditional characteristics). We formulate a scaling law metric(*n*) = *E* + *A n*^−*α*^ and evaluate it with five metrics: accuracy (MAE, RMSE), distributional calibration (KS distance, goodness-of-fit), and training objective (negative log loss) using a neural temporal point process trained on a cohort of four-ECG datasets. The law fits all five metrics. While point accuracy is near saturation at *n* = 183, KS distance and goodness-of-fit improve by 6% and 12% respectively when extrapolated to 10,000 subjects, showing that scaling decisions in stochastic domains must be guided by distributional calibration rather than point accuracy.

## 1 Introduction

Scaling laws turn data budgets into a quantitative decision: by fitting a power law between held-out loss and data size, we can accurately determine training budget [1]. Such laws presume the target is predictable: given enough context, the next token or label can be determined. For heartbeat intervals, even a perfect model of the conditional intensity cannot predict the next interval exactly [2]. A scaling curve fit to point error (MAE, RMSE) flattens early due to irreducible error from beat to beat variability, but this is not indicative of true saturation. While accuracy evaluates the error of individual data points, distributional calibration (measured by Kolmogorov-Smirnov distance and goodness-of-fit) assess whether predicted probability densities match true conditional distributions. Given that clinical data collection is expensive and subject to regulation and privacy concerns, determining the right cohort size requires evaluating scaling by both distributional calibration and point accuracy. This principle extends across stochastic domains, including traffic, seismicity, and system latency telemetry, where the generating process can be modeled by a known family (e.g., point processes).

## 2 Methods

We model heartbeat intervals *r*_*i*_ = *t*_*i*_ − *t*_*i*−1_ with a density-based neural temporal point process [3, 4]. Log-normalized intervals pass through a GRU history encoder, and a linear projection maps the hidden state to the parameters of a *K*=8 component log-normal mixture for *p*(*r*_*i*+1_ | ℋ_*i*_), the next interval given the history ℋ_*i*_ = {*r*_1_, …, *r*_*i*_}. We pool 229 subjects from four datasets ([5],[6],[7]) and representatively sample ≈80%-10%-10% for training, validation and testing.

We evaluate scaling across five metrics. To measure distributional calibration (whether our predicted distributions match the true distributional characteristics), we apply the time-rescaling theorem [8, 4], calculating **Kolmogorov-Smirnov (KS) distance** against a uniform distribution. From this, we define **goodness-of-fit** as the fraction of 2-minute windows with a KS distance below the 5% significance cutoff 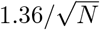. We also report point accuracy metrics **beat-wise MAE and RMSE**, plus training objective **negative log loss(NLL)**.

Models are trained over ten random seeds at each *n* ∈ {1, 2, 4, 8, 16, 32, 48, 64, 96, 128, 160, 183}, with seed-level (cluster) bootstrapping. We fit a power law metric(*n*) = *E* + *A n*^−*α*^, where *E* is the irreducible floor (theoretic minimum error even with infinite data), *A* is the magnitude of initial error due to data scarcity, and *α* is the data-efficiency exponent that measures how performance improves as data size increases. We define metric saturation as the point where the reducible error component *A n*^−*α*^ falls within 1% of the irreducible floor *E*. The law is fit in log space for all metrics except NLL (which is fit linearly as it is negative) using a robust Huber loss. We report estimated parameters with 95% CIs, and the projected metric values when scaling the data further.

## 3 Results

The proposed law holds across metrics (Table 1, Figure 1); fits for KS distance and goodness-of-fit are well constrained (*R*^2^ = 0.86, 0.90), while the fits for MAE and RMSE are noisier (*R*^2^ = 0.51, 0.45) due to irreducible beat-to-beat variability. Extrapolating to 10,000 subjects (Table 2) shows that MAE and RMSE are near saturation at *n* = 183, while KS distance improves by 6%, and goodness-of-fit improves from 86.9% to 97.5%.

**Table 1:** Fitted scaling law coefficients (with 95% CIs). ^*^Fit as deficit to imposed 100% ceiling. ^†^*R*^2^ is log-space except for NLL, which is fit in linear space.

| Metric | $E$ | $A$ | $\alpha$ | $R^{2\dagger}$ |
| --- | --- | --- | --- | --- |
| KS distance | 0.0776 (0.068, 0.086) | 0.35 (0.27, 0.47) | 0.80 (0.64, 1.05) | 0.86 |
| goodness-of-fit* | 100 (imposed) | 110.7 (100.9, 121.5) | 0.41 (0.37, 0.44) | 0.90 |
| MAE (ms) | 49.6 (44.9, 54.9) | 53.2 (30.0, 77.6) | 1.01 (0.58, 1.41) | 0.51 |
| RMSE (ms) | 71.1 (65.5, 79.7) | 53.3 (36.5, 77.7) | 0.77 (0.52, 1.09) | 0.45 |
| NLL (nats/beat) | -1.940 (-2.034, -1.848) | 1.771 (1.465, 2.271) | 0.835 (0.638, 1.143) | 0.479 |

**Table 2:** Projections from the fitted laws.

| Subjects $n$ | KS dist. | goodness-of-fit | MAE (ms) | RMSE (ms) | NLL (nats/beat) |
| --- | --- | --- | --- | --- | --- |
| 183 (pooled cohort) | 0.0829 | 86.9 | 49.85 | 72.07 | −1.9168 |
| 1,000 | 0.0790 | 93.5 | 49.62 | 71.36 | −1.9341 |
| 10,000 | 0.0778 | 97.5 | 49.58 | 71.14 | −1.9389 |
| $n \rightarrow \infty$ | 0.0776 | 100 (imposed) | 49.57 | 71.10 | −1.9397 |

**Figure 1:**
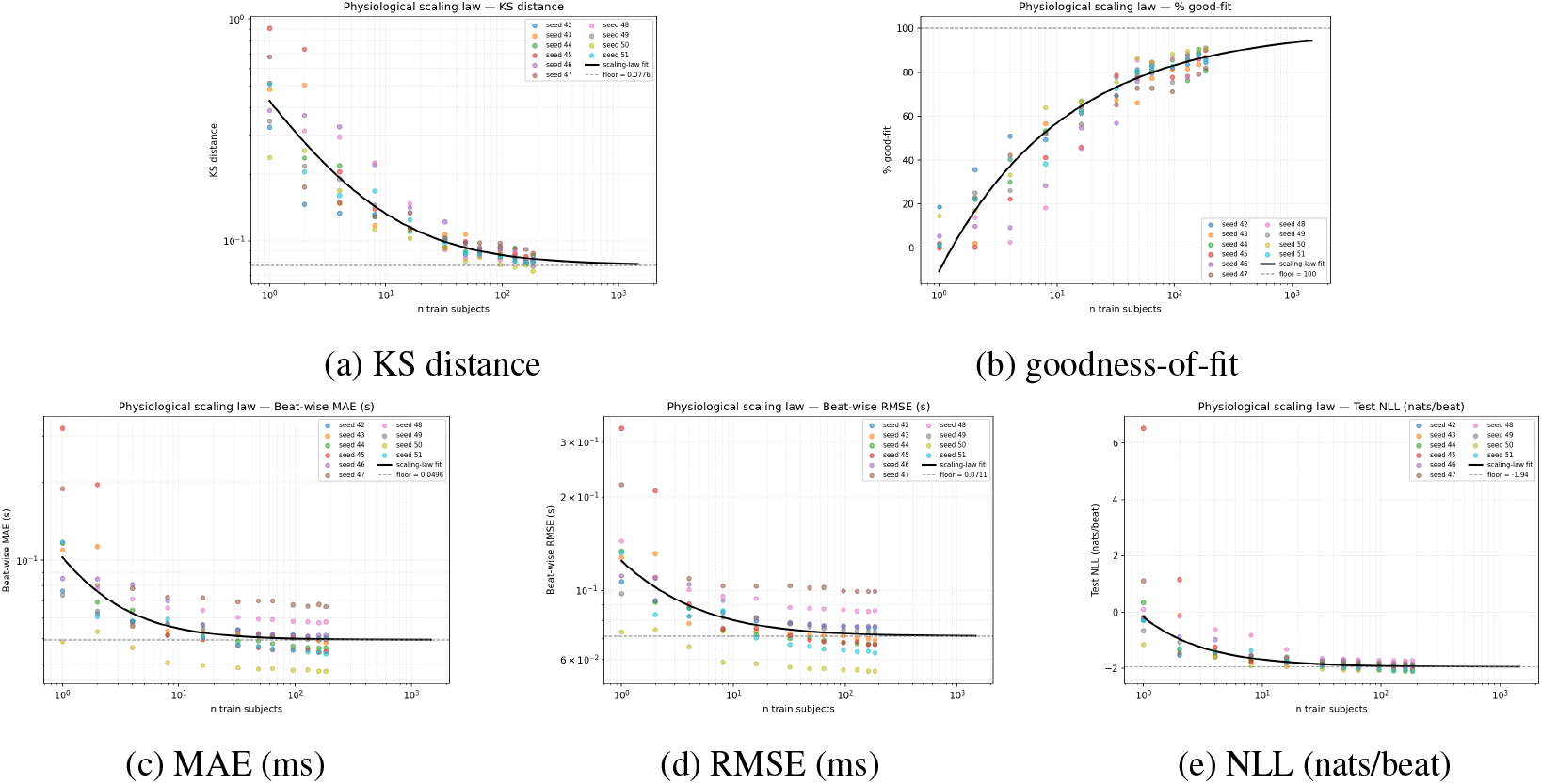
Fitted scaling laws vs. number of training subjects (axes are log-log except goodness-of-fit and NLL, which use a linear y-axis). Points are 10 seeds× 12 subset sizes; solid curves are metric(*n*) = *E* + *A n*^−*α*^, dashed lines the fitted floors *E*.

## 4 Discussion

Standard ML scaling work defines E as a residual parameter that absorbs unexplained variance, but in our work it quantifies the beat-to-beat variability that additional data cannot reduce. Evaluating scaling with point accuracy incorrectly shows that performance is near saturation at *n* = 183. Distributional calibration (measured by KS distance and goodness-of-fit), continues to improve from 183 to 10,000 subjects. In stochastic domains where the underlying process can be explicitly modeled (e.g., hazard rates, seismicity, etc.), relying on point accuracy alone results in under-scaled models, and we should use distributional calibration to guide data budget decisions. Future work can extend this framework to joint parameter-data scaling in physiology, or other modeled stochastic systems.

